# Synthetic lethal interaction between WEE1 and PKMYT1 is a target for multiple low dose treatment of high-grade serous ovarian carcinoma

**DOI:** 10.1101/2023.03.31.535053

**Authors:** Jan Benada, Daria Bulanova, Violette Azzoni, Valdemaras Petrosius, Saba Ghazanfar, Krister Wennerberg, Claus Storgaard Sørensen

## Abstract

Ovarian cancer is driven by genetic alterations that necessitate protective DNA damage and replication stress responses through cell cycle control and genome maintenance. This creates specific vulnerabilities that may be exploited therapeutically. WEE1 kinase is a key cell cycle control kinase, and it has emerged as a promising cancer therapy target. However, adverse effects have limited its clinical progress, especially, when tested in combination with chemotherapies. A strong genetic interaction between WEE1 and PKMYT1 led us to hypothesize that a multiple low dose approach utilising joint WEE1 and PKMYT1 inhibition would allow exploitation of the synthetic lethality. In the present study, we found that the combination of WEE1 and PKMYT1 inhibition exhibited synergistic effects in eradicating ovarian cancer cells and ovarian organoid models at a low dose. The WEE1 and PKMYT1 inhibition synergistically promoted activation of CDK1 by decreasing the phosphorylation levels of T14 and Y15 residues. Furthermore, the combined treatment exacerbated DNA replication stress and replication catastrophe, leading to increase of the genomic instability and inflammatory STAT1 signalling activation. Finally, the multiple low dosing was well tolerated in mice. These findings suggest a new multiple low dose approach to harness the potency of WEE1 inhibition through the synthetic lethal interaction with PKMYT1 that may contribute to the development of new treatments for ovarian cancer.

## Introduction

High-grade serous carcinoma (HGSC) is both the most prevalent and most fatal subtype of ovarian cancer. Standard therapy for HGSC comprise of cytoreductive surgery, followed by chemotherapy with DNA-damaging agents, such as platinum drugs, either alone or in combination with taxane drugs *(1*–*3)*. While primary tumours usually respond favourably to the treatment, over 80% of cases relapse and more than half of these acquire resistance to the treatment. Moreover, the relapsed HGSC is typically fast-growing and invasive *(1*–*3)*. Platinum-resistant tumours are subjected to salvage therapy with DNA-damaging drugs including doxorubicin, topotecan, etoposide, vinorelbine and gemcitabine. However, the response rate at this stage is only about 10–15% with a median progression-free survival of 3–4 months, underscoring the urgent need for better treatments *(1*–*3)*.

A proposed therapeutic option for HGSC is the use of WEE1 inhibitors *(4)*. WEE1 catalyses inhibitory phosphorylation on Tyrosine 15 of both CDK1 and CDK2 and thus limits the CDK activity *(5, 6)*. WEE1 inhibition deregulates CDK activity and exacerbates DNA replication stress to intolerable levels, effectively killing the cells in the process termed replication catastrophe *(7, 8)*. Mechanistically, unrestricted CDK activity leads to excessive replication origin firing, which in turn results in depletion of protective RPA. RPA exhaustion marks a point of no return, when unprotected replicons collapse in lethal genome-wide DNA breakage *(7, 9)*. Moreover, WEE1 inhibition also overrides the G2/M DNA damage checkpoint forcing cells to enter mitosis with unrepaired DNA which triggers cell death *(10)*. Stringent G2/M checkpoint is especially vital for cancer cells, as they frequently lose the ability to arrest their cell cycle and repair DNA in the G1 checkpoint *(11, 12)*.

The most studied inhibitor of WEE1 is adavosertib (AZD1775, MK1775), which has been the focal point of multitude clinical studies *(13)* (clinicaltrials.gov). In particular, positive outcomes for HGSC were reported in phase II clinical trials by combining adavosertib and gemcitabine treatment, exploiting high levels of replication stress in HGSC (4). However, despite the years of testing, adavosertib has not reached clinical use, which is mainly due to significant adverse effects when used in combination with chemotherapy *(14, 15)*.Of note, several other WEE1 inhibitors are currently evaluated in clinical trials, including ZN-c3 (Zentalis), Debio0123 (Debiopharm), IMP7068 (Impact Therapeutics) and SY4835 (Shouyao Holding) (clinicaltrials.gov).

An attractive strategy to limit unacceptable toxicity is to use a lower dose of inhibitors but at same time target multiple proteins of a single signalling pathway to still achieve a complete pathway inhibition. This multiple low dose therapy also reduces cancer selective pressure against a single target that may result in treatment resistance *(16*–*18)*. A prime candidate for synergistic effect with WEE1 is the kinase PKMYT1. PKMYT1 phosphorylates CDK1 at Threonine 14 and thus inhibits CDK activity *(19)*. WEE1 and PKMYT1 have been described to be synthetically lethal in CRISPR-Cas9 screens in glioblastoma stem-like cells *(20)*. Moreover, upregulation of PKMYT1 has been shown to promote resistance of cancer cells to WEE1 inhibition *(21)*.

PKMYT1 as an anticancer target is much less studied than WEE1, however, Repare Therapeutics have recently identified a first-in-class PKMYT1 inhibitor, RP-6306 (22). RP-6306, used as single compound, showed promising *in vitro* results for ovarian cancer cells *(23)*, and is currently being fast tracked to clinical trials (clinicaltrials.gov; NCT05147350, NCT05147272, NCT04855656, NCT05605509, NCT05601440). Notably, PKMYT1 expression was reported to be upregulated in ovarian cancers and correlated with poor prognosis, making it an appealing therapeutic target *(24)*. The availability of a PKMYT1 inhibitor prompted us to investigate the synergistic potential of its combined application with WEE1 inhibition.

## Results

### Combined inhibition of WEE1 and PKMYT1 synergize in killing of cancer cells

WEE1 inhibition has emerged as a strategy to eliminate cancer cells, however, adverse effect concerns have warranted further preclinical investigations. We noted that combined WEE1 and PKMYT1 genetic ablation was lethal in a glioma setting (20). To evaluate the potential synergistic effect of cotargeting WEE1 and PKMYT1, we conducted a dose response matrix for cell viability with the WEE1 inhibitor adavosertib in combination with the PKMYT1 inhibitor RP-6306 (Fig.1A and B). The U2OS cell line was selected as model since it has been well characterized for WEE1 and CDK functions, and notably, investigated in detail for the effects of adavosertib treatment (25). In the 100 nM concentrations the combinatorial treatment led to efficient killing of most cancer cells. In stark contrast, treatment with 100 nM of single compound treatment had negligible effect on cell viability (Fig.1B). A synergy analysis revealed a strong synergistic interaction between the inhibitors (Fig.1C). Furthermore, these findings were reproduced using the combination of RP-6306 with another clinically relevant WEE1 inhibitor, ZN-c3 (Suppl.FigA and B). The efficacy of used inhibitors was validated by Western blots, assaying the substrates of WEE1 and PKMYT1 - CDK1pY15 and CDK1pT14, respectively (Suppl.Fig.1C and D).

**Fig.1.**
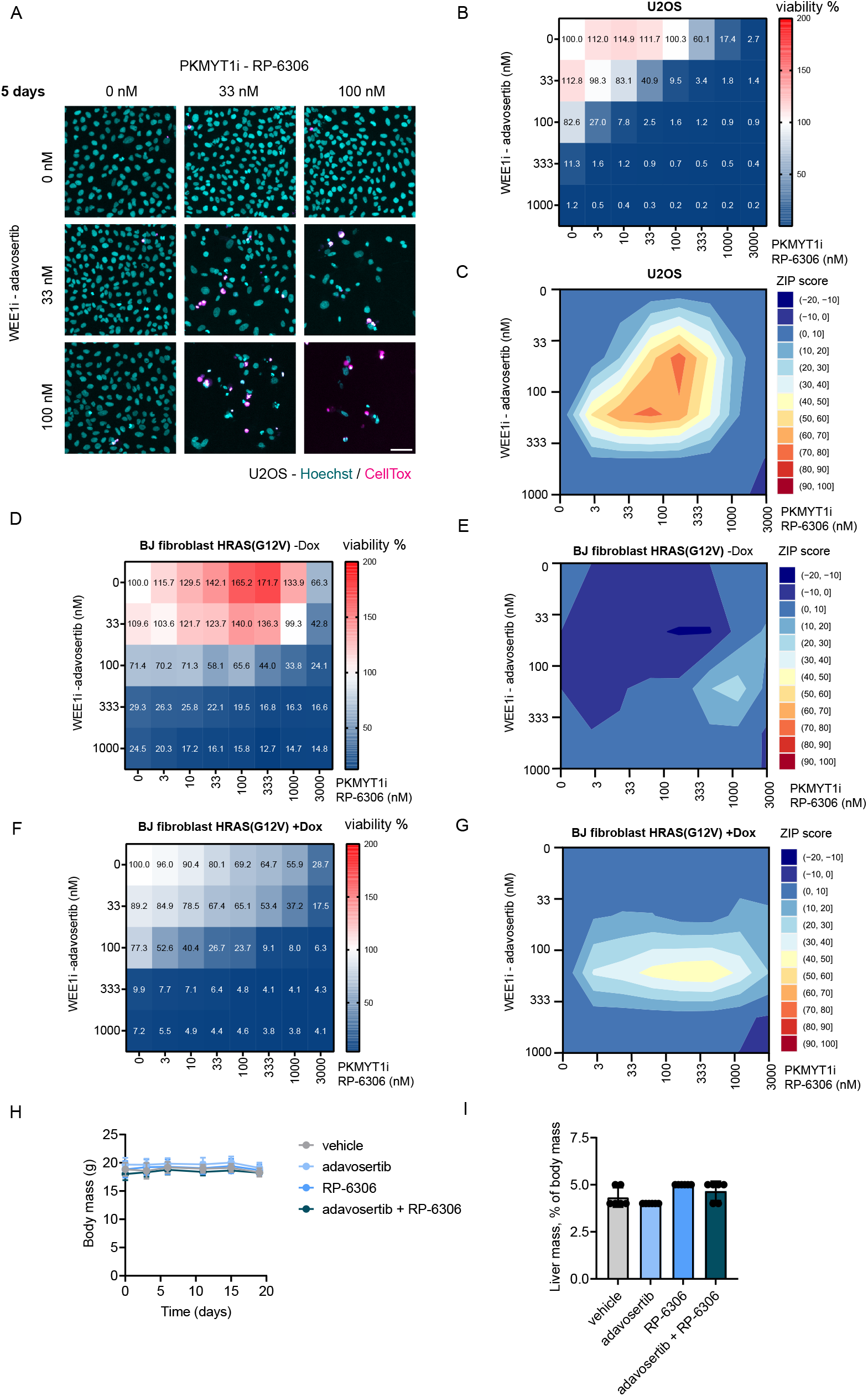
Combined inhibition of WEE1 and PKMYT1 synergize in killing of cancer cells: **(A)** Representative images of cell viability assay upon 5-day treatment with adavosertib in combination with RP-6306 in U2OS cells; scale bar represents 50 μm **(B)** Dose response matrix for cell viability upon 5-day treatment with adavosertib in combination with RP-6306 in U2OS cells, data represent mean from triplicate. **(C)** Synergy ZIP scores corresponding to data in (B) presented a synergy landscape. A score ≥ 10 represents synergy, a score ≤ −10 represents antagonism. **(D)** Dose response matrix for cell viability upon 5-day treatment with adavosertib in combination with RP-6306 in BJ fibroblast HRAS(G12V) Tet ON cells without doxycycline, data represent mean from triplicate. **(E)** Synergy ZIP scores corresponding to data in (D) presented a synergy landscape. **(F)** Dose response matrix for cell viability upon 5-day treatment with adavosertib in combination with RP-6306 in BJ fibroblast HRAS(G12V) Tet ON cells with doxycycline induced expression, doxycycline has been added 2 days before inhibitor treatment and together with inhibitors, data represent mean from triplicate. **(G)** Synergy ZIP scores corresponding to data in (F) **(H)** Changes in body weight in NGX mice treated with RP-6306 (5 mg/kg) adavosertib (15 mg/kg), the combination thereof, or vehicle. Drugs were administered orally twice daily for 21 days at an intermittent dosing schedule. Results are expressed as mean body mass ± SD, n = 6 for each cohort. **(I)** Liver weight normalized to body weight at the end of the treatment described in (H). Bars represent mean ± SD.

In contrast to normal cells, cancer cells are characterized by multiple genetic alterations in driver genes that promote high rate of proliferation and impose replication stress (RS) *(26, 27)*. The high level of RS renders cancer cells particularly dependent on safeguarding mechanisms and these can be exploited by cancer treatments. To assess whether the multiple low dose application of WEE1i and PKMYT1i preferentially eliminated high RS cells and less so their normal counterparts, we conducted dose response matrix in normal BJ fibroblast cells with doxycycline inducible oncogene HRAS(G12V) *(28, 29)*. To first confirm the impact of HRAS(G12V) induction, we monitored proliferation rates which indeed were elevated by activating HRAS signalling (Suppl.Fig.1E). Moreover, normal non-induced BJ fibroblast were indeed more resistant to the treatment and exhibited cell death only in higher doses compared to U2OS cells (Fig.1D). This was particularly marked for the response to the PKMYT1 inhibitor. Accordingly, the synergy score was comparably lower in the non-induced setting, and the inhibitors synergized only in a very limited concentration window (Fig.1E). Importantly, HRAS(G12V) induction sensitised the BJ cells to the treatment (Fig.1F and G). To further investigate the impact of combinatorial treatment on normal cells, we administered adavosertib and RP-6306 by oral gavage to mice. The achieved plasma levels (when corrected to plasma protein binding) of both compounds were in concentration range where we observed synergistic effect in killing cancer cells *in vitro* (Suppl.Fig.F-H). Notably, compounds had little impact on mouse bodyweight and liver size when administered individually or jointly (Fig.1H and I). This suggests that when administered at low doses, the combinatorial treatment is tolerated in mouse models.

### WEE1i and PKMYT1i co-inhibition exacerbates replication stress and triggers replication catastrophe

Next, we aimed to characterise in detail the mechanism of action of the WEE1i and PKMYT1i combination. WEE1 and PKMYT1 suppress replication stress, and they also guard against premature mitotic entry even in cells experiencing genotoxic challenges through replication stress *(10, 23)*. Thus, we reasoned that combined WEE1i and PKMYT1i could elevate replication stress to intolerable level. To assess replication stress levels, we employed quantitative imagebased cytometry (QIBC) to measure accumulation of ssDNA by quantifying the levels of chromatin-bound RPA and phosphorylation of histone H2AX on serine 139 (γH2AX) as a marker of DNA damage *(7)*. The dose response matrix of acute treatment (4 h) displayed a synergistic effect of adavosertib and RP-6306 in inducing the replication stress, which in higher inhibitor concentration propagated into replication catastrophe (RC) (Fig.2A and B). We observed the same impact for the combination of ZN-c3 and RP-6306 (Suppl.Fig.2A and B). Given the role of WEE1 and PKMYT1 as master regulators of CDK1, we reasoned that induction of replication stress and catastrophe corresponded to increased levels of CDK activity in S-phase. Indeed, we observed that CDK1 activity, measured as phosphorylation of the CDK substrate FOXM1 at threonine 600 *(30)*, increased rapidly upon the combined treatment and correlated with induction of replication stress as measured by QIBC (Fig.2C-E). Moreover, we were able to reverse the replication catastrophe phenotype by inhibition of CDK activity with CDK1i RO-3066 (Fig.2C-E) *(31)*. Complementary to QIBC, western blot analysis of cellular fractionates showed increased CDK activity, as measured by pan-CDK substrate, and increased chromatin loading of RPA which indicates replication stress. We also observed increased phosphorylation and activation of markers of the DNA damage response such as CHK1 phosphorylated at serine 345, CHK2 phosphorylated at threonine 68, and γH2AX (Fig.2F). Collectively, the data indicated a marked replication stress and replication catastrophe response to combined WEE1 and PKMYT1 inhibition that likely explain the major treatment lethality in cancer cells. Moreover, cells which do not die in imminent replication catastrophe will be forced by high CDK activity into premature mitotic entry with high levels of replication stress-generated DNA damage resulting in further loss of cell fitness.

**Fig.2.**
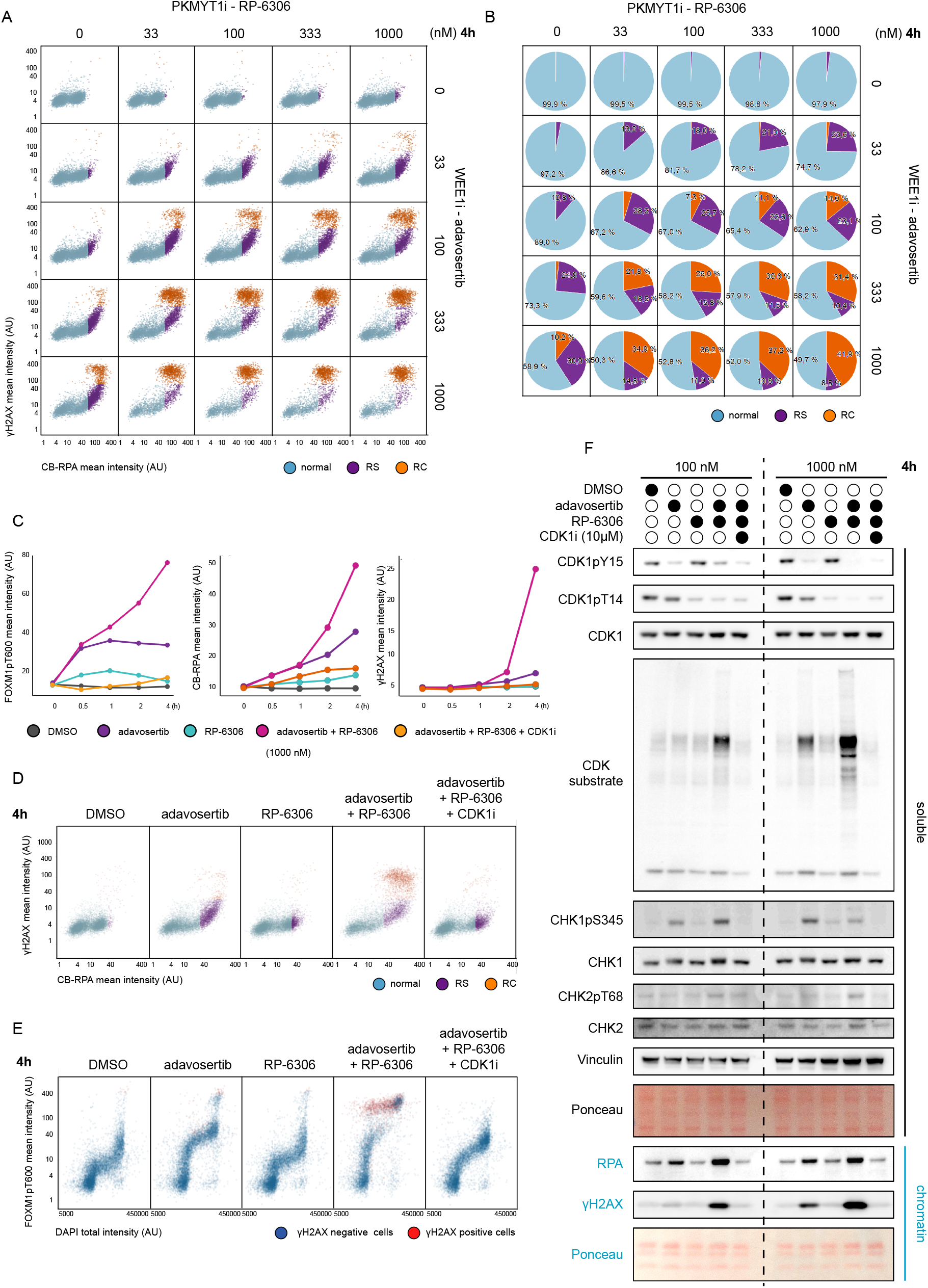
WEE1 and PKMYT1 co-inhibition exacerbates replication stress and triggers replication catastrophe: **(A)** Dose response matrix for QIBC analysis of replication stress upon 4 h treatment with adavosertib in combination with RP-6306 in U2OS cells. AU = arbitrary unit, RS = replication stress, RC = replication catastrophe **(B)** Analysis of relative cell populations percentage from (A). **(C)** Line plots for CDK activity (FOXM1pT600), chromatin bound RPA (CB-RPA) and γH2AX in indicated treatments. **(D)** QIBC plots related to (C) for CB-RPA and γH2AX. **(E)** QIBC plots related to (C) for FOXM1pT600 and DAPI, red indicate γH2AX-positive cells; AU = arbitrary unit. **(F)** Western blot analysis of chromatin fractionation in indicated treatments; CDK1i = RO-3306.

### Combined WEE1 and PKMYT1 inhibition increases genomic instability and activates a cGAS-STING response

Increased levels of replication stress have been associated with exacerbation of genome instability and formation of micronuclei after mitotic progression with DNA damage *(32, 33)*. To test micronuclei formation in our system, we treated the U2OS cells with combinatorial low-dose WEE1i and PKMYT1i for 3 days and assessed the percentage of cells with micronuclei. We observed a significant increase in their formation with combinatorial WEE1i and PKMYT1i treatment in doses as low as 33 nM (Fig.3A and B). The increased presence of micronuclei is linked to activation of innate immunity and the clearance of tumours *in vivo (32)*. Mechanistically, this is mediated by the cGAS-STING pathway and subsequent activation of STAT signalling response. Accordingly, we observed increased accumulation of cGAS (Fig.3A and Fig.3C) and elevated marker of STAT1 activation (STAT1pY701) (Fig.3D). Taken together these data demonstrate that the WEE1i and PKMYT1i combination activates cGAS-STING response.

**Fig.3.**
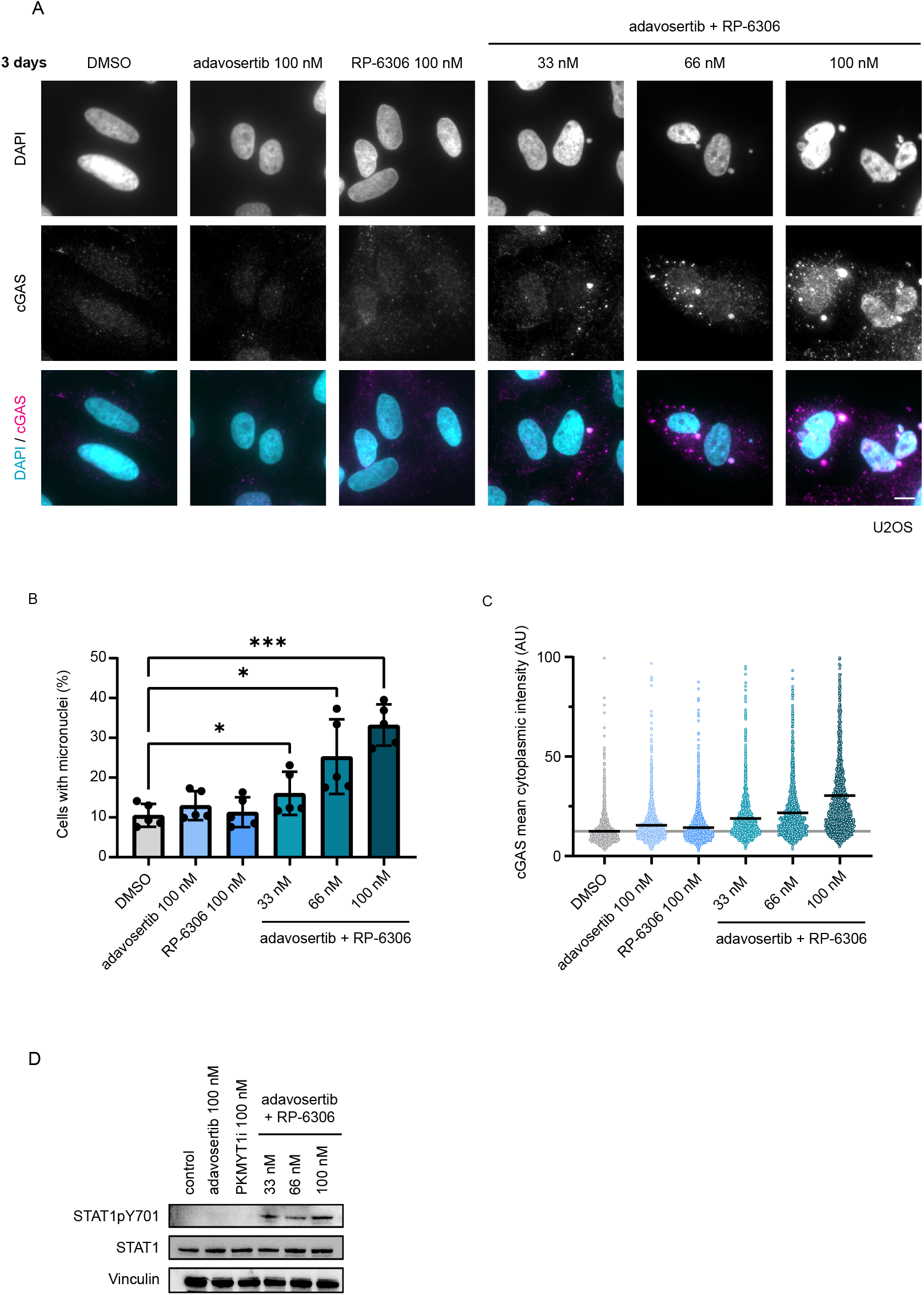
WEE1 and PKMYT1 co-inhibition increases genomic instability and activates cGAS-STING response: **(A)** Representative images of micronuclei formation and cGAS activation upon 3-day treatment with adavosertib in combination with RP-6306 in U2OS cells; scale bar represents 10 μm. **(B)** Quantification of micronuclei formation activation upon 3-day treatment with adavosertib in combination with RP-6306 in U2OS cells; bars indicate mean and SD from biological quintuplicate; two-way analysis of variance (ANOVA), *0.01<P≤0.05, **0.001<P≤0.01, ***0.0001<P≤0.001 **(C)** Quantification of cytoplasmic cGAS intensity upon 3 day treatment with adavosertib in combination with RP-6306 in U2OS cells, bars indicate mean. **(D)** Western blot analysis of STAT1 activation (STAT1pY701) upon 72 h treatment with adavosertib in combination with RP-6306 in U2OS cells.

### WEE1i and PKMYT1i multiple low dose treatment is efficient against a variety of HGSC cell lines regardless of driver oncogene

As described above, combined WEE1i and PKMYT1i application might be suited for aggressive hard-to-treat cancers with high proliferation rates such as HGSC. To address this, we tested a diverse panel of HGSC cell lines in the dose response matrix for cell viability. These included OVCAR3, OVCAR8, COV318, COV362 and KURAMOCHI cell lines. We previously profiled these cell lines for their expression of driver oncogenes (Suppl.Fig.3A) *(34)*. Regardless of the expression of specific driver oncogenes, all ovarian cancer cell lines were eradicated by WEE1i and PKMYT1i following multiple low dose exposure (Fig.4A-J). Upon tailored inspection and like U2OS, we observed replication stress and induction of replication catastrophe in OVCAR3 (high cyclin E; Suppl.Fig.3B and C) and KURAMOCHI (high KRAS; Suppl.Fig.3D and E) after the WEE1 and PKMYT1 in multiple-low dose treatment. We also noted that all tested ovarian cancer cell lines responded in a similar dose range to U2OS, with COV318 displaying a slightly different pattern. COV318 represents a highly heterogenous cell line and the majority of COV318 population responded with even higher sensitivity than the other HGSC cell lines, though approximately 10% of the cells survived the treatment.

**Fig.4.**
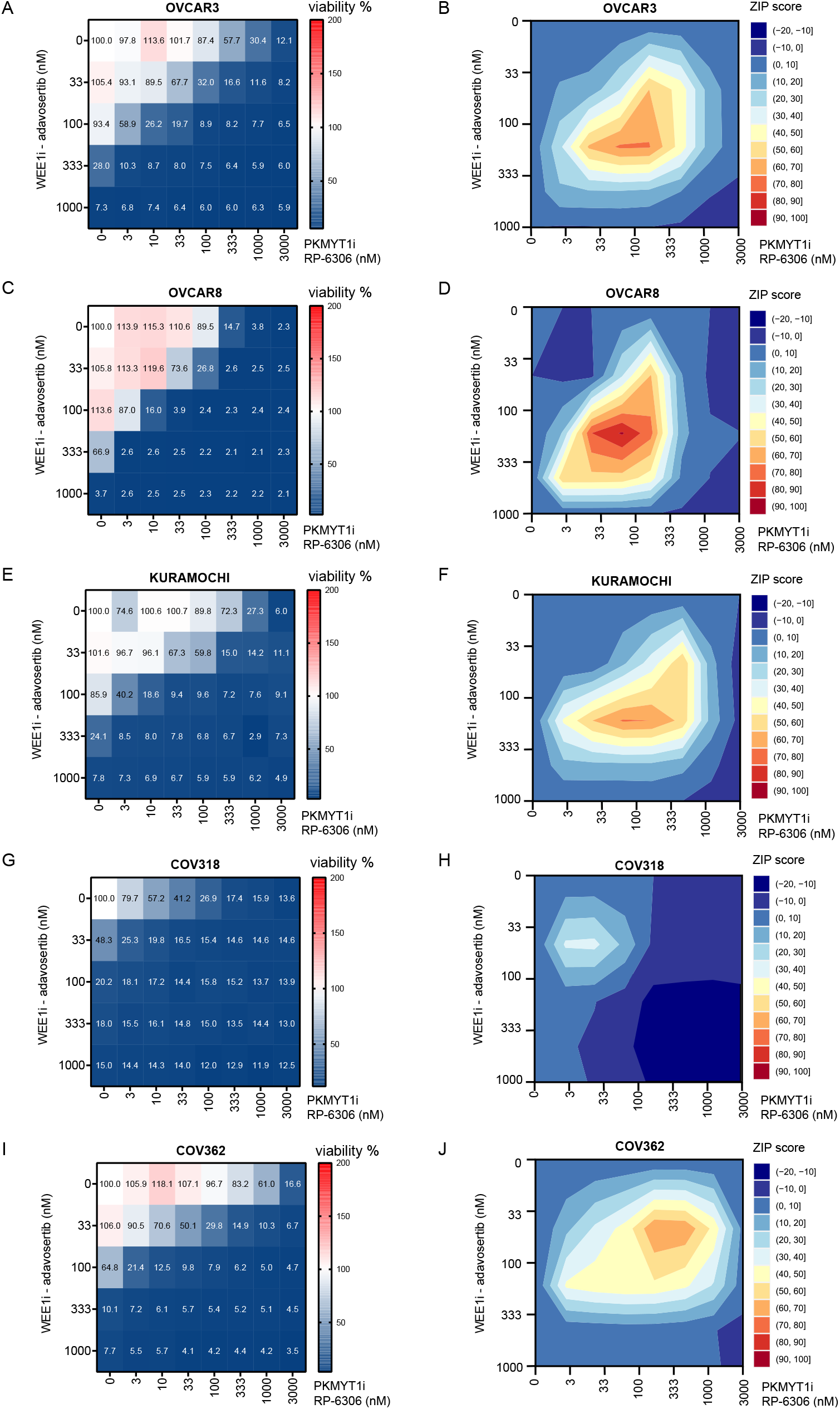
WEE1 and PKMYT1 co-inhibition kills a variety of high-grade ovarian serous adenocarcinoma cell lines regardless of their driver oncogenes: **(A)** Dose response matrix for cell viability upon 5-day treatment with adavosertib in combination with RP-6306 in OVCAR3 cells, data represent mean from triplicate. **(B)** Synergy ZIP scores corresponding to data in (A) presented a synergy landscape. A score ≥ 10 represents synergy, a score ≤ −10 represents antagonism. **(C)** Dose response matrix for cell viability upon 5-day treatment with adavosertib in combination with RP-6306 in OVCAR8 cells, data represent mean from triplicate. **(D)** Synergy ZIP scores corresponding to data in (C) presented a synergy landscape. **(E)** Dose response matrix for cell viability upon 5-day treatment with adavosertib in combination with RP-6306 in KURAMOCHI cells, data represent mean from triplicate. **(F)** Synergy ZIP scores corresponding to data in (E) presented a synergy landscape. A score ≥ 10 represents synergy, a score ≤ −10 represents antagonism. **(G)** Dose response matrix for cell viability upon 5-day treatment with adavosertib in combination with RP-6306 in COV318 cells, data represent mean from triplicate. **(H)** Synergy ZIP scores corresponding to data in (G) presented a synergy landscape. **(I)** Dose response matrix for cell viability upon 5-day treatment with WEE1i inhibitor adavosertib in combination with PKMYT1 inhibitor RP-6306 in COV362 cells, data represent mean from triplicate. **(J)** Synergy ZIP scores corresponding to data in (I) presented a synergy landscape.

### WEE1i and PKMYT1i multiple low dose approach eradicates patient-derived ovarian cancer organoids

To further assess the potential for translation of the WEE1i and PKMYT1i multiple low dose approach, we evaluated its efficacy in a set of clinically relevant HGSC patient-derived organoid cultures that retained genetic makeup and heterogeneity of the tumor of origin (Summarized in Fig5A) *(35)*.Imaging-based toxicity assay revealed dose-dependent synergistic efficacy of the combination in all tested HGSC organoid cultures (Fig.5B-F), independent of CCNE1, MYC or KRAS amplification, the site of origin of the tumor cells, or previous exposure to the replication stress-inducing carboplatin-based neoadjuvant chemotherapy (NACT). Importantly, organoid cultures EOC989 and EOC884, that were derived from residual tumour cells from patients treated with chemotherapy (NACT), showed prominent synergistic response to the combined WEE1 and PKMYT1 inhibition. Both WEE1 and PKMYT1 inhibitors are tested in clinical trials in combination with gemcitabine (clinicaltrials.gov, NCT02101775, NCT05147272). Therefore, we also addressed potential impact of combining gemcitabine with our multiple low dose approach. We pre-treated select ovarian cancer cells (OVCAR3 and KURAMOCHI) and organoids (EOC884 and EOC989) with gemcitabine for 18 h followed by combined WEE1 and PKMYT1 inhibition. We observed a considerable impact as cells were more readily eradicated following gemcitabine pre-treatment (Suppl.Fig.4A-D). This suggests potential in combining standard chemotherapy with the WEE1i and PKMYT1i multiple low dose approach.

**Fig.5.**
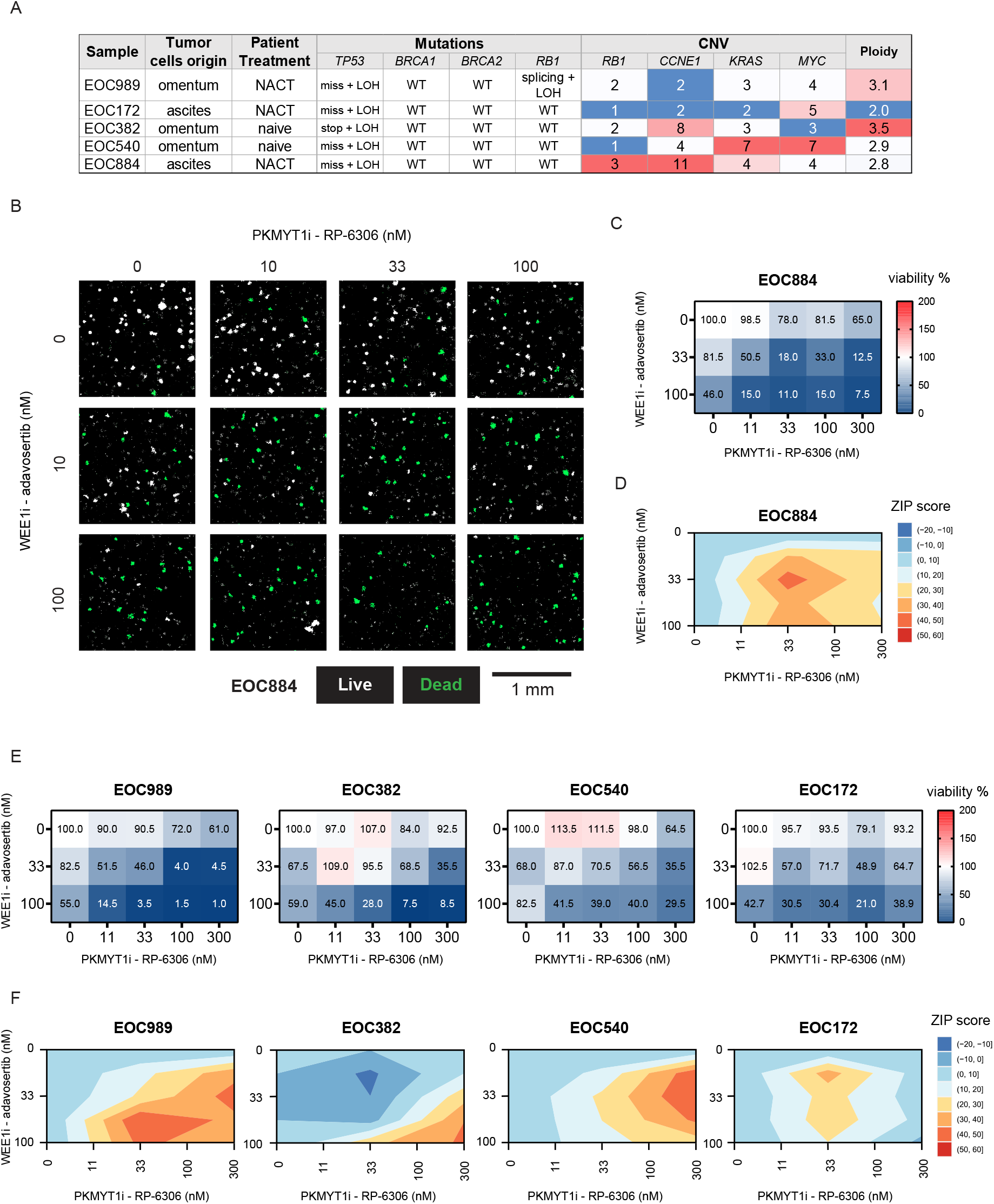
WEE1 and PKMYT1 co-inhibition kills patient-derived ovarian cancer organoids: **(A)** Characteristics of the selected set of long-term organoid cultures. NACT, neoadjuvant platinum-based chemotherapy. Mutations: fs, frameshift; stop, stopgain; miss, missense mutation; splicing, splicing isoform mutation; LOH, loss of heterozygosity. Copy number columns indicate number of gene copies per cell, the numbers in red highlight the amplification of the gene. **(B)** Representative micrographs of HGSC organoids from patient EOC884. The organoids were treated with increasing doses of RP-6306 and adavosertib for 7 days. White masks label CellTox Green-negative live organoids, green masks label the CellTox Green-positive dead organoids. Scale bar indicates 1 mm **(C)** Quantification of the viability of EOC884 organoids after 7 days of the drug combination treatment illustrated in (B). Values in heat map cells present the percentage of live organoids relative to DMSO-treated control. Mean, n=3. **(D)** ZIP synergy plots for adavosertib and RP-6306 interaction in the dose matrix viability testing in EOC884 organoids in (C). Interaction was classified as synergistic if ZIP score ≥ 10 **(E)** Quantification of the viability in panel of HGSC organoids after 7 days of the drug combination treatment illustrated in B. Values in heat map cells present the percentage of live organoids relative to DMSO-treated control. Mean, n=3. **(F)** ZIP synergy plots for adavosertib and RP-6306 interaction in the dose matrix viability testing in panel of HGSC (E). Interaction was classified as synergistic if ZIP score ≥ 10.

## Discussion

Here, we report the combinatorial drugging of PKMYT1 and WEE1 kinases, which synergistically eradicates cancer cells already at low drug dose. Our data highlight the potential for multiple low dose treatment and support the notion that combining full dose treatments may not be the only approach when administrating a set of targeted drugs. Synthetically lethal interactions are attractive in cancer treatment as they may allow reduced adverse effects while drugging cancerspecific vulnerabilities in a similar manner as the targeting of BRCA deficiency with PARP inhibition *(36)*. However, few drug candidates and treatments are currently based on synthetic lethality and the multiple low dose approach is still under development. The multiple low dose treatment may represent a desirable strategy also for targeting other checkpoint kinases as Golder at al., 2022 demonstrated that multiple low dose combinatorial drugging of ATR and CHK1 inhibitors proved effective in killing HGSC cells *(37)*. Our data show that cancer cells or oncogene exposed cells are more sensitive than normal cells to the WEE1i and PKMYT1i combinatorial treatment, which suggests the evolution of sensitivity during cancer development. It was recently demonstrated that PKMYT1 inhibition with RP-6306 displays synthetic lethality with CCNE1 amplification based on the marked replication stress induced by CCNE1 overexpression *(23)*. Indeed, cancer cells generally display elevated replication stress due to activated drivers such as CCNE1, KRAS and MYC *(26, 27)*. We observed that oncogenic HRAS(G12V) expression also sensitized cells to PKMYT1 inhibition alone and to combined WEE1i and PKMYT1i treatment. In addition, combined WEE1i and PKMYT1i treatment was effective in killing a diverse panel of HGSC cell lines and organoids, regardless of their driver oncogene. This suggests that the treatment efficacy is not limited to a particular oncogene, but rather to a more general feature of oncogene-induced replication stress. In support of this notion, we have observed an exacerbation of replication stress and induction of replication catastrophe, which likely mechanistically underlay a major part of combinatorial treatment lethality in cancer cells. Moreover, cell cycle control in cancer cells is perturbed by the frequent hits in the p53 and RB pathways *(38)*. This creates a dependency on the remaining cell cycle control mechanisms and causes cancer cell vulnerabilities to targeted treatments that interfere with these mechanisms. Thus, increased replication stress and limited cell cycle control mechanisms make cancer cells reliant on G2 phase control to limit detrimental premature mitotic entry. Deregulated passage through G2-M transition promotes mitotic cell death and it also triggers micronuclei formations, which in turn leads to innate immunity activation such as cGAS-STING mediated interferon responses *(32, 33, 39)*. In agreement with this finding, we observed induced cGAS-STING signalling at low doses, suggesting that the multiple low dose approach may yield additional benefits *in vivo* by triggering the immune cell-mediated tumour clearance. We also observed that the low dose combination was well tolerated in mouse models, although additional studies are needed to evaluate the *in vivo* efficacy of the low dose combination, as well as tolerability in higher species.

The ubiquitous TP53 mutations in HGSC *(40)* as well as alterations in G1/S transition master regulators (RB1 deficiency *(41)*, CCNE1 amplifications *(42)*) present the highly relevant molecular landscape for exploration of WEE1 and PKMYT1 co-inhibition as a treatment option. It is of interest to assess the potential of the combination treatment to eradicate the residual tumour cells, which lose the sensitivity to the standard chemotherapeutics *(43)* and frequently are selected to restore the HR repair pathway *(44)*, excluding the further possibility to use PARP inhibitors in the treatment. We have employed ovarian cancer organoids to represent three-dimensional cell culture models closely reflecting the primary tissue’s biology and pathology. We observed a pronounced synergistic response to the combination in HGSC organoids established from residual tumour samples collected after a few cycles of chemotherapy (EOC884, EOC989, Figure 5, Suppl.Fig.4C and D). As the preclinical drug testing in organoids helps accurately predict the clinical treatment outcome *(35, 45*–*47)*, the observed efficacy of the multiple low dose treatment with WEE1 and PKMYT inhibitors suggests that the respective tumours may have responded to the therapeutic combination. Hence, the combination may offer a treatment strategy to attenuate the relapses due to chemoresistant disease, which affects up to 70% of HGSC cases *(1, 3)*. Even though we focused mostly on ovarian cancer model systems, we have also shown that the multiple low dose strategy can potentially be effective in other tumour types as indicated by our results in U2OS cells. Collectively, our findings argue for a reduced focus on maximal tolerable dose of single targeted cancer drugs and suggest the use of drug combinations at low and non-toxic concentrations.

## Material and Methods

### Cell lines

Human osteosarcoma cell line U2OS (ATCC) and COV362 (ATCC) were cultured in Dulbecco’s Modified Eagle’s medium (DMEM; Gibco) supplemented with 10% foetal bovine serum (FBS; Cytiva) and 1% Penicillin-Streptomycin (10,000 U/mL; Gibco). High-grade serous ovarian cancer cell lines OVCAR3, OVCAR8 (NCI Tumor Repository (Frederick, MD) and KURAMOCHI (Japanese Collection of Research Bioresources Cell Bank (JCRB)) cell lines were cultured in Roswell Park Memorial Institute 1640 medium (RPMI1-1640, Gibco) supplemented with 10% foetal bovine serum and 1% Penicillin-Streptomycin. BJ fibroblast HRAS(G12V) Tet ON cells were cultured in Dulbecco’s Modified Eagle’s medium (DMEM; Gibco) supplemented with 10% Tet-System Approved foetal bovine serum (FBS; Gibco) and 1% Penicillin-Streptomycin (10,000 U/mL; Gibco). HRAS(G12V) expression was induced by 2 μg/mL of doxycycline. All cells were cultured at 37°C with 5% CO2 and checked for mycoplasma infection regularly.

### Organoid cultures

Long-term HGSC organoid cultures were established and characterized earlier as described (35). The samples were cultured in 7.5 mg/mL BME-2 matrix (Cultrex, BioTechne) in sample-specific media (35) - for EOC883, EOC172 Medium 1 (Advanced DMEM/F12 (#12634010, Gibco), supplemented with 100 μg/mL Primocin (#ant-pm-1, Invivogen), 10 mM HEPES (#15630080, Gibco), 1 mM N-acetylcysteine (#A7250, Sigma), 1X GlutaMAX (#35050061), 1X B-27 Supplement, 0.5 μM SB202190 (#HY-10295, MedChemExpress), 0.5 μM A83-01 (#SML0788, Sigma), 10 ng/mL recombinant human FGF-10 (#100-26, Peprotech), 10 ng/mL recombinant human FGF-4 (#100-31, Peprotech), 100 nM β-estradiol (#E2758, Sigma), 5 mM nicotinamide (#N0636, Sigma.); for EOC989, EOC540, EOC382 Medium 2 (Medium 1 supplemented with 5 ng/mL EGF, 5 μM heregulin-1β, 0.5 μg/mL hydrocortisone and 5 μM forskolin).

### Organoid drug sensitivity assay

The organoids were dissociated in Tryple Express solution (Thermo Fisher) as described previously *(35)*. Dissociated cells were resuspended in 7.5 mg/mL BME-2 matrix gel at 2-5×104 cells/mL and seeded in 10 μL droplets to individual wells of 96-well Cell Carrier Ultra plates (Perkin Elmer). Once settled, 200 μL the organoid-specific growth medium containing 5 μM ROCK inhibitor Y-27632 (HY-10583, MedChemExpress) was added. After 4 days, the growth medium was changed to 200 μL of medium containing adavosertib or RP-6306 at the indicated concentrations, or 25 μM staurosporine as a positive control for cytotoxicity. After 7 days, organoids were stained using Hoechst 33342 (1 μg/mL) and CellTox Green (Promega, 1/20 000) dyes for 8 h prior to imaging at Opera Phenix (Perkin Elmer) confocal screening microscope. The fraction of dead organoids was discriminated by the CellTox Green signal per organoid from the confocal images analysis using Harmony software (Perkin Elmer).

### Drug sensitivity assay

Drugs diluted in DMSO to the desired concentrations were dispensed at 30 nL volume to 384-well black plates (Corning, cat. 3864) using an Echo 550 acoustic liquid handler (Labcyte). Cell-killing benzethonium chloride (BzCl, 100 μM) and compound vehicle (DMSO, 0.1%) were used as positive and negative controls, respectively. Cells were diluted to medium at the desired number per mL and the suspension was dispensed to the pre-drugged plates at 30 μL. Alternatively, drugs were dispensed by manual pipetting into 96 well plates (Greiner-BIO) and cells were dispensed at the desired number of 100 μL. After 5 days of incubation at 37°C, 10 μL/30 μL for 384-well/96-well plates, respectively, of PBS containing 4 μg/mL Hoechst 33342 and 1/10 000 CellTox Green dyes were added for 1 h prior to imaging. Images were obtained automatically with the ScanR acquisition software controlling a motorized Olympus IX-83 wide-field microscope, equipped with Lumencor SpectraX light engine and Hamamatsu ORCA-FLASH 4.0, using Olympus Universal Plan Super Apo 4 × / 0.16 AIR Objective.

### Quantitative image-based cytometry

Cells growing on either 96-well microplates (Greiner-BIO) or 12 mm coverslips were treated with different combinations of drugs for indicated time intervals. After the treatment, the media was quickly removed and the cells were incubated in pre-extraction buffer (25 mM HEPES, pH 7.5, 50 mM NaCl, 1 mM EDTA, 3 mM MgCl2, 300 mM sucrose, and 0.5% Triton X-100) on ice for 2 min and immediately fixed in formaldehyde 4% (VWR) for 10 min at the room temperature. For the analysis of the micronuclei number and cGAS the pre-extraction step was omitted. Primary antibodies (γH2AX 1:300, Cell Signaling Technology, Cat#2577; RPA 1:300, Millipore, Cat#MABE285; FOXM1pT600 1:1000, Cell Signaling Technology, Cat#14655; cGAS 1:300, Cell Signaling Technology, Cat#15102) were diluted in filtered DMEM containing 10% FBS and 5% Bovine Serum Albumin (BSA; Sigma). Incubations with the primary antibodies were performed at RT for 1 h. Microplates were washed three times with 0.05% PBS-Tween20 and incubated in DMEM/FBS/BSA containing secondary fluorescently labelled antibodies (Alexa Fluor dyes (1:1000; Thermo Fisher Scientific) and DAPI (0.5 mg/mL; Sigma-Aldrich) for 1 h at room temperature. Images were obtained automatically with the ScanR acquisition software controlling a motorized Olympus IX-83 wide-field microscope, equipped with a Lumencor SpectraX light engine and Hamamatsu ORCAFLASH 4.0. Olympus PlanC N 10 × / 0.25 AIR objective was used to capture γH2AX, RPA and QIBC data. Micronuclei images were obtained with a 0.75 AIR UPlanSApo 40 × / 0.95 AIR objective. Images were processed and quantified using the ScanR image analysis software for total nuclear pixel intensities for DAPI (Arbitrary units: A.U.) and mean (total pixel intensities divided by nuclear area) nuclear intensities (A.U.) for γH2AX, chromatin-bound RPA and FOXM1pT600. Micronuclei were segmented based on DAPI channel within a cytoplasmic mask surrounding the nucleus. Similarly, cGAS intensity was determined within the cytoplasmic mask. Further analysis and data visualisation was then carried out with Tibco Spotfire software (Tibco, RRID: SCR_008858). Representative images were processed using ImageJ/Fiji (RRID:SCR_002285, imagej.net).

### Western blotting

Cells were lysed in RIPA (Sigma) buffer containing EDTA-free protease inhibitor cocktail (Roche) and phosphatase inhibitors (Roche). Lysates were treated with Benzonase Nuclease (Sigma-Aldrich) for 30 min on ice. Lysates were centrifuged for 15 min at 20 000 × g at 4°C. Protein concentration was then measured with Bradford assay and adjusted accordingly to ensure equal loading. Lysates were mixed with 4 × Laemmli sample buffer (Sigma) and boiled for 10 min at 95°C. Samples were run on NuPAGE Bis-Tris 4-12% gels according to manufacturer instructions. Proteins were then transferred to a nitrocellulose membrane and blocked with PBS + 0.1% Tween 20 + 5% Milk powder (Sigma) and incubated overnight with primary antibodies at 4°C. The membrane was then washed 3 × 5 min in PBS + 0.1% Tween 20 and incubated with secondary HRP conjugated antibodies for 2 h at room temperature. Membranes were again washed 3 × 5 min with PBS + 0.1% Tween 20 and incubated with Classico/Crescendo Western HRP substrate (Millipore-Sigma) for 2 min. Chemiluminescence signal was detected using a Bio-Rad ChemiDoc Touch Imaging System. Following primary antibodies were used: STAT1pY701 1:100, Cell Signaling Technology, Cat#9171; STAT1 1:1000, Cell Signaling Technology, Cat#9176; Vinculin 1:10000, Sigma, Cat#V9131; CDK1 1:1000, Abcam, Cat#ab18, CDK1pT14 1:1000, Abcam, Cat#ab58509; CDK1pT15 1:1000, Cell Signaling Technology, Cat#9111S; phospho-CDK substrate motif 1:1000, Cell Signaling Technology, Cat#9477; CHK2 1:500, Santa Cruz, Cat#sc-56296; CHK2pT18 1:500, Cell Signaling Technology, Ca 2661; CHK1 1:500, Santa Cruz, Cat#sc-56291; CHK2pS345 1:500, Cell Signaling Technology, Cat#2348; γH2AX 1:1000, Cell Signaling Technology, Cat#2577; RPA70 1:1000, Abcam, Cat#ab79398.

### Cell fractionation

Cells were grown in 10 cm dishes, treated as indicated, washed three times with ice-cold PBS and harvested. The soluble fractions were extracted by incubation in ice-cold nuclear buffer (10 mM HEPES pH 7, 200 mM NaCl, 1 mM EDTA, 0.5% NP-40) supplemented with protease and phosphatase inhibitors (Roche) for 10 min on ice, and centrifuged at 2000 × g for 6 min. The remaining pellet was rinsed once with ice-cold washing buffer (10 mM HEPES pH 7, 50 mM NaCl, 0.3 M sucrose, 0.5% Triton X-100) supplemented with protease and phosphatase inhibitors (Roche), which was removed by centrifugation at 1400 × g for 6 min. Chromatin fractions were extracted by incubation in RIPA buffer (50 mM Tris-HCl pH 8, 150 mM NaCl, 1% IGEPAL CA-630, 0.1% SDS, 0.1% Na-deoxycholate) supplemented with protease and phosphatase inhibitors (Roche) and Benzonase Nuclease (Sigma) for 30 min on ice and clarified by centrifugation at maximum speed.

### Drug-drug interaction analysis

To assess the outcome of the drug combination treatment, we applied the zero interaction potency (ZIP) synergy model *(48)*. The ZIP score reflects the additional cell line response induced by the combinatorial treatment compared to the expected response based on the two single compounds. A ZIP score ≥ 10 is considered synergistic, a score ≤ −10 represents antagonism. ZIP scores were calculated for each combination in the dose matrix by SynergyFinder 2.0 *(49)*; (synergyfinder.org/ and synergyfinder.fimm.fi)and plotted as synergy landscapes using RStudio (RRID:SCR_000432, www.r-project.org) and ggplot2 package (ggplot2.tidyverse.org)(*50)*.

### Statistical analysis

Statistical analyses were conducted using GraphPad Prism (v.9.5.1). For multiple comparisons, statistical significance (adjusted P values) was calculated using the two-way analysis of variance (ANOVA), Tukey multiple comparisons test, Welch ANOVA test with Dunnetts multiple comparisons test or unpaired Student’s t test. Results are reported as non-significant at P>0.05, and with increasing degrees of significance symbolized by the number of asterisks: *: 0.01<P≤0.05, **: 0.001<P≤0.01, ***: 0.0001<P≤0.001 and ****: P≤0.0001. Statistical details for each experiment can be found in the corresponding legend.

### In vivo drug tolerance study

All experiments were carried out under authorization and guidance from the Danish Inspectorate for Animal Experimentation. Female 7-week-old mice of NGX (NOD-Prkdc scid-IL2rg Tm1/Rj) strain (Janvier Labs) were randomized in 4 treatment cohorts of 6 animals. The mice were housed in individually ventilated cages with a humidity of 55%±10%, a temperature of 22±2°C, and a dark/light cycle of 12h/12h with light from 6:00 to 18:00. adavosertib (Repare Therapeutics) and RP-6306 (Repare Therapeutics) were formulated in 1% DMSO, 0.5% methylcellulose and administered by oral gavage at 15 mg/kg and 5 mg/kg, respectively, alone or in combination. For intermittent 21-day dosing, the drugs, the combination, or the vehicle were given twice daily, with 8 h interval, for 5 days a week, followed by 2 treatment-free days. Animal weight was measured twice weekly, and the overall animal condition was monitored daily. At the end of the treatment, all mice were humanely sacrificed, and liver weights were measured.

### Pharmacokinetics

Whole blood samples were collected 30 min and 8 h after the first drug treatment. Immediately after collection, 20 μL of blood were mixed with 60 μL of 0.1 M citrate buffer (0.1 M trisodium citrate, pH 7.4) and stored at −80°C before the analysis. All samples were quantified using a reversed-phase liquid chromatography gradient coupled to electrospray mass spectrometry operated in positive mode. PK parameters were calculated using non-compartmental analysis.

## Supporting information

Manuscript and figures 1-5

## ACKNOWLEDGEMENTS

We thank the members of CSS and KW laboratories and Repare Therapeutics for insightful comments; Repare Therapeutics for providing us with RP-6306; Shou Yun Yin (Repare Therapeutics) for performing the pharmacokinetic analysis; Department of Experimental Medicine (University of Copenhagen) for animal work; the core facilities at BRIC for assistance, BRIC’s Core Facility for High-Content CRISPR Screens and Karolin Voßgröne for assistance with Echo 550 acoustic liquid handler; J. Bartek laboratory (Danish Cancer Society) for BJ fibroblast HRAS (G12V) Tet ON cells. Funding: Supported by the Danish Cancer Society and the Novo Nordisk Foundation. The manuscript was formatted in Overleaf (www.overleaf.com) using the modified “HenriquesLab bioRxiv” template by Ricardo Henriques.

