## Supplementary material for "Synthetic lethal interaction between WEE1 and PKMYT1 is a target for multiple low dose treatment of high-grade serous ovarian carcinoma": Manuscript and figures 1-5

### Supplementary Materials

Supplementary Figures 1-4

preprint

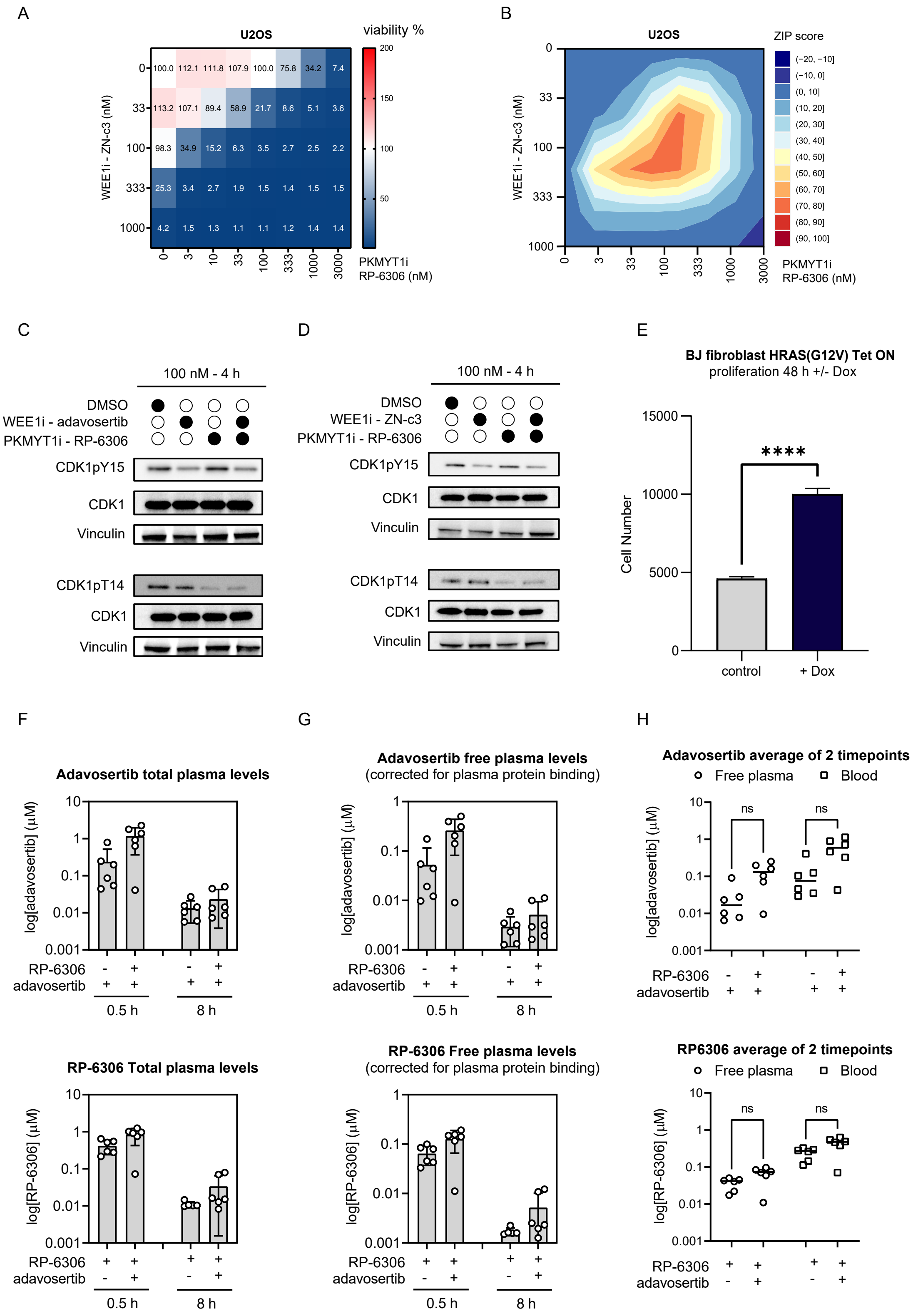


**Suppl.Fig.1: (A)** Dose response matrix for cell viability upon 5-day treatment with WEE1i inhibitor ZN-c3 in combination with RP-6306 in U2OS cells, data represent mean from triplicate. **(B)** Synergy ZIP scores corresponding to data in (A) presented a synergy landscape. A score $\geq$ 10 represent synergy, a score $\leq$ -10 represents antagonism. **(C)** Western blot analysis of CDK1pY15 and CDK1pT14 upon 4 h treatment either with adavosertib or RP-6306 or in combination in U2OS cells. **(D)** Western blot analysis of CDK1pY15 and CDK1pT14 upon 4 h treatment either with WEE1i inhibitor ZN-c3, RP-6306, or in combination in U2OS cells. **(E)** Proliferation of BJ fibroblast HRAS(G12V) Tet ON without or with doxycycline addition for 48 h, bars indicate mean and SD, n = 25, ****: P < 0.0001 (unpaired Student's t test). **(F)** Total plasma levels of RP-6306 and adavosertib related to Fig.1.H and G, n = 6, bars indicate mean and SD **(G)** Free plasma levels (corrected for plasma protein binding) of RP-6306 and adavosertib related to Fig.1.H and G, n = 6, bars indicate mean and SD. **(H)** The average total and free plasma levels of the 2 timepoints presented in (F) nad (G), Kruskal-Wallis test.

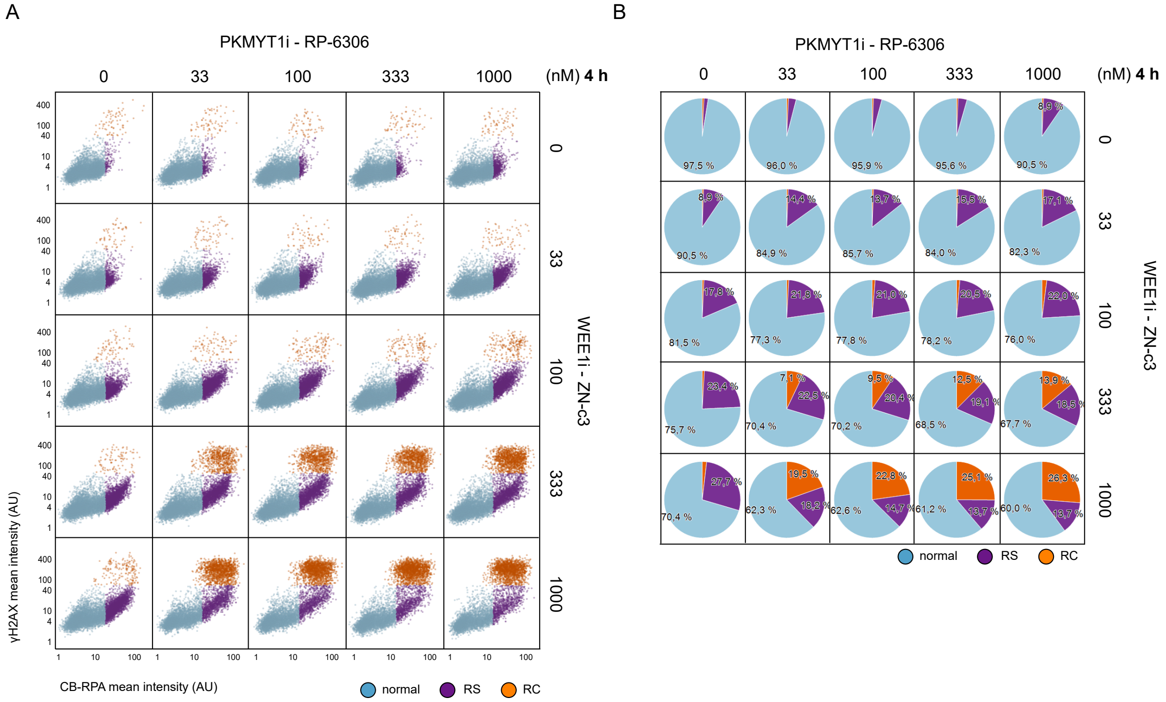


**Suppl.Fig.2: (A)** Dose response matrix for QIBC analysis of replication stress upon 4 h treatment with WEE1i inhibitor ZN-c3 in combination with RP-6306 in U2OS cells. AU = arbitrary unit, RS = replication stress, RC = replication catastrophe **(B)** Analysis of relative cell populations percentage from (A) RS = replication stress, RC = replication catastrophe.

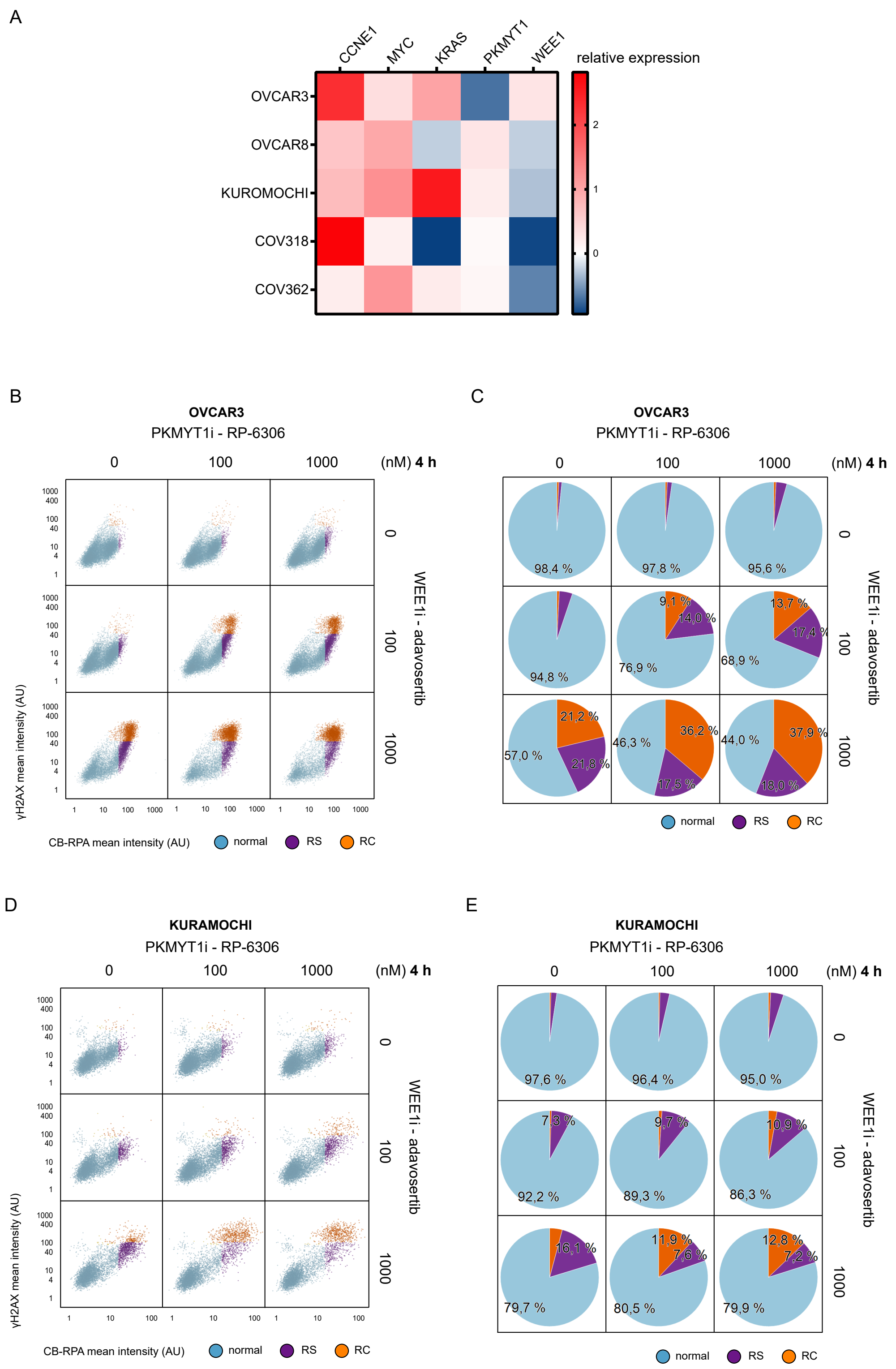


**Suppl.Fig.3: (A)** Expression of selected oncogenes, WEE1 and PKMYT1 for panel of HGSC cell lines. **(B)** Dose response matrix for QIBC analysis of replication stress with adavosertib in combination with RP-6306 in OVCAR3 cells. AU = arbitrary unit, RS = replication stress, RC = replication catastrophe **(C)** Analysis of relative cell populations percentage from (B) **(D)** Dose response matrix for QIBC analysis of replication stress with adavosertib in combination with PKMYT1 inhibitor RP-6306 in KURAMOCHI cells. **(E)** Analysis of relative cell populations percentage from (D).

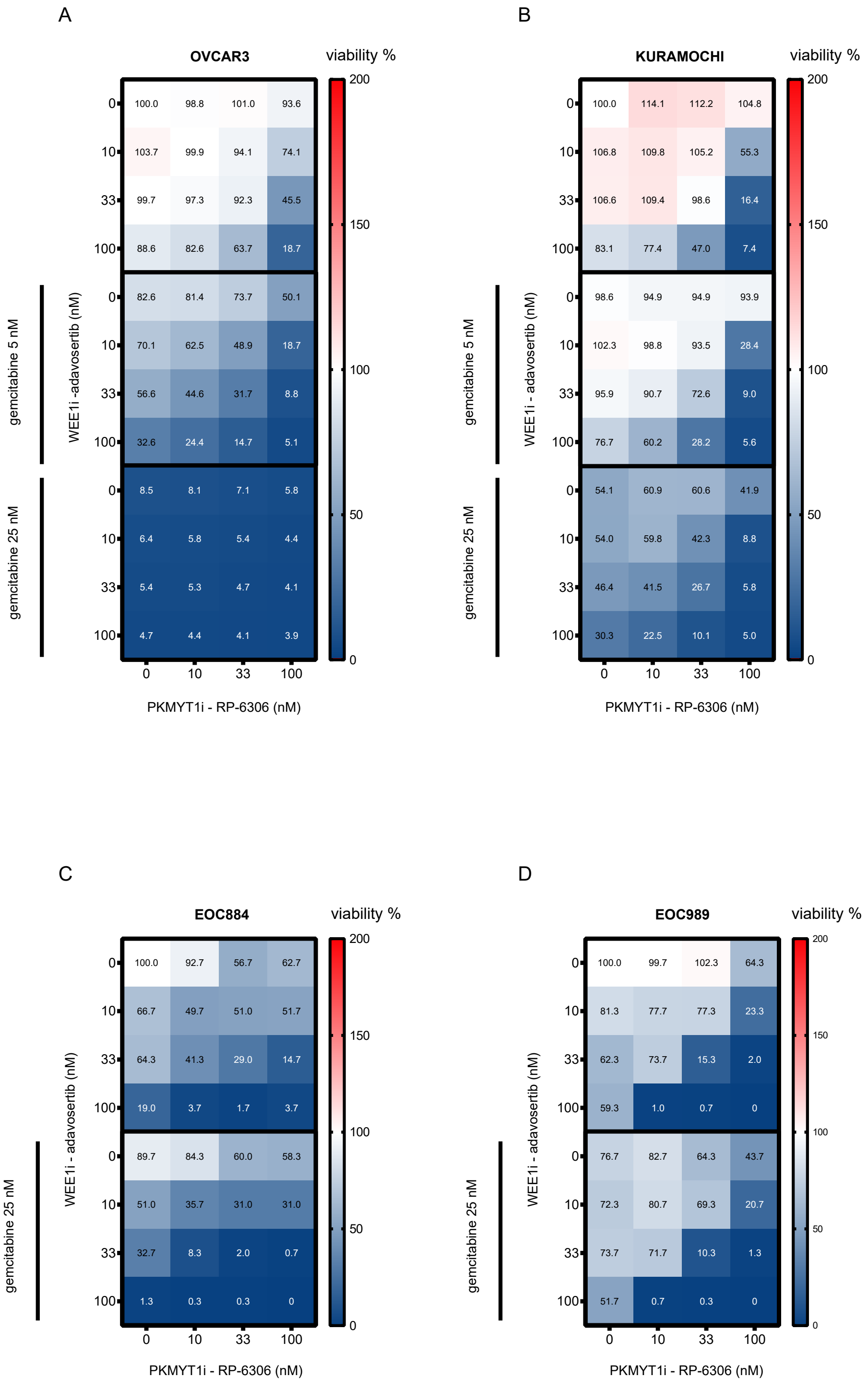


**Suppl.Fig.4: (A)** Dose response matrix for OVCAR3 cells viability upon 5-day treatment with WEE1i inhibitor adavosertib in combination with PKMYT1 inhibitor RP-6306 without and with 18h pre-treatment with 5 nM or 25 nM gemcitabine, respectively; mean, n= 3. **(B)** Dose response matrix for KURAMOCHI cells viability upon 5-day treatment with WEE1i inhibitor adavosertib in combination with PKMYT1 inhibitor RP-6306 without and with 18 h pre-treatment with 5 nM or 25 nM gemcitabine, respectively; mean, n = 3. **(C)** Dose response matrix for EOC884 organoids viability upon 7 day treatment with WEE1i inhibitor adavosertib in combination with PKMYT1 inhibitor RP-6306 without and with 18 h pre-treatment with 25 nM g emcitabine, data represent percentage of live organoids relative to DMSO-treated control, mean, n = 3 **(D)** Dose response matrix for EOC989 organoids viability upon 7-day treatment with WEE1i inhibitor adavosertib in combination with PKMYT1 inhibitor RP-6306 without and with 18 h pre-treatment with 25 nM gemcitabine, data represent percentage of live organoids relative to DMSO-treated control, mean, n = 3.
